# A Yap-dependent transcriptional program directs cell migration for embryo axis assembly

**DOI:** 10.1101/2022.02.06.479280

**Authors:** Ana Sousa-Ortega, Javier Vazquez-Marin, Estefanía Sanabria-Reinoso, Rocío Polvillo, Alejandro Campoy-López, Lorena Buono, Felix Loosli, María Almuedo-Castillo, Juan R. Martinez-Morales

**Author notes:** Both authors contributed equally to this work (Ana Sousa-Ortega, Javier Vazquez-Marin).

## Abstract

The condensation of the embryo primary axis is a fundamental landmark in the establishment of the vertebrate body plan. Although the complex morphogenetic movements directing cell convergence towards the midline have been described extensively, little is known on how gastrulating cells interpret mechanical cues. Yap proteins are among the best characterized transcriptional mechanotransducers, yet their role in gastrulation has remained elusive. Here we show that the double knockout of *yap* and its paralog *yap1b* in medaka results in an axis assembly failure. Quantitative live imaging reveals that mutant cells display reduced displacement and migratory persistence. By characterizing the Yap-dependent transcriptional program, we identified genes involved in cytoskeletal organization and cell-ECM adhesion, rather than in germ layer specification, as direct Yap targets. Dynamic analysis of Tead sensors and downstream targets reveals Yap is acting in migratory cells, and not as a midline beacon, to direct gastrulating precursors trajectories by promoting cortical actin recruitment and focal adhesions assembly. We propose that Yap is engaged in a mechano-regulatory loop that is essential to maintain the directed cell migration sustaining embryo axis formation.

## INTRODUCTION

The assembly of the primary embryo axis is an essential event for the foundation of the body plan in all bilaterian animals. During gastrulation, the embryo axis emerges through evolutionary conserved morphogenetic movements, which operate in coordination with the linage restriction events that lead to the specification of the three basic germ layers; ectoderm, mesoderm and endoderm (Keller, 2002; Solnica-Krezel and Sepich, 2012). Basic gastrulation movements, such as emboly, epiboly, convergence and extension, follow a similar logic in all species, particularly within the same animal phyla. In fact, the capacity to form and elongate an axis is apparently an intrinsic property of the developing cell collectives. Thus, even in the absence of stereotypic embryo cues (e.g. extraembryonic tissues and embryo geometry), isolated ESCs populations have the capacity to self-organize into a rudimentary A-P axis *in vitro*, as demonstrated for mammalian gastruloids (Beccari et al., 2018; Moris et al., 2020) and zebrafish blastoderm aggregates (Fulton et al., 2020; Schauer et al., 2020). Despite commonalities, the regulatory weight of specific cellular and molecular mechanisms behind general gastrulation rearrangements may vary among species, reflecting diverse embryo size, geometry, and adaptations to ecological niches (Kalinka and Tomancak, 2012). An example of these context-dependent adaptations comes from the comparative analysis of convergent-extension cellular behaviors between xenopus and zebrafish. Whereas mediolateral intercalation plays an important role in axis development in both vertebrates, dorsally directed migration has a relevant contribution only in teleosts (Wallingford et al., 2002). It is likely that, by harnessing conserved developmental modules to specific embryo adaptations, these context-dependent mechanisms are essential to confer robustness to the gastrulation process. Morphogenetic robustness, as it is observed in developing embryos, is indeed one of the main features that distinguish gastrulation *in vivo* from self-organized morphogenesis *in vitro* (Anlas and Trivedi, 2021; Schauer and Heisenberg, 2021). Mechanical cues stand up among the candidate contextual inputs that may act as channeling mechanisms to maintain gastrulation as an invariant process. Mechano-regulatory loops have an essential function in maintaining homeostasis during development and tissue remodeling (Hannezo and Heisenberg, 2019), being dysregulated mechanical feedbacks a common landmark in numerous pathologies, particularly in cancer (Northcott et al., 2018). In the context of gastrulation, it has been shown that, mechanical strains play a conserved role in mesoderm specification both in Drosophila and zebrafish through nuclear translocation of the transcriptional regulator ß-catenin (Brunet et al., 2013). This seems to be an ancestral mechano-regulatory module as it has been also reported in Cnidarian embryos (Pukhlyakova et al., 2018). Despite, their relevance, the impact of mechanotransduction and mechanosensation mechanisms on gastrulation dynamics has been explored only recently (e.g.(Chanet et al., 2017; Mitrossilis et al., 2017; Schwayer et al., 2019).

Yap proteins are well-known transcriptional regulators able to shuttle to the nucleus upon mechanical stimulation, being active in cells that have undergone cell spreading and inactive in round and compact cells (Dupont et al., 2011). Their ability to sense mechanical strains depends on acto-myosin contractility, actin capping and severing proteins, the integrin-talin mechanosensitive clutch, as well as a direct coupling between the extracellular matrix and the nuclear envelope (Aragona et al., 2013; Elosegui-Artola et al., 2017; Elosegui-Artola et al., 2016). Initially characterized as effectors of the Hippo signaling cascade, Yap proteins play a key role both during embryogenesis, as master regulators of growth, cell specification, and survival (Varelas, 2014), as well as in adult organs, where they are critical for tissue repair and cancer progression (Zanconato et al., 2016). More recently an increasing number of reports have linked Yap proteins to cell rearrangements and tissue morphogenesis (Davis and Tapon, 2019; Porazinski et al., 2015). This is not unsurprising, given Yap/Taz transcriptional ability to modulate cytoskeletal and extracellular matrix (ECM) components, as reported in mammalian cell lines (Calvo et al., 2013; Nardone et al., 2017). A shortened and widened A-P axis has been repeatedly observed in *yap1* deficient embryos from different vertebrate models including mouse, (Morin-Kensicki et al., 2006), xenopus (Gee et al., 2011), zebrafish (Kimelman et al., 2017) and medaka (Vazquez-Marin et al., 2019); which may suggest a conserved role in convergent-extension. However, despite these observations, the mechanistic link between Yap proteins and gastrulation movements has remained elusive.

Here we show that the simultaneous inactivation of *yap1* and *yap1b*, the two members of the Yap family in medaka (Vazquez-Marin et al., 2019), results in a complete failure to assemble the posterior half of the embryo axis. The analysis of the transcriptional program activated by Yap proteins indicates that, with the noticeable exception of the non-neural ectoderm, basic germ layers specification does not depend on Yap function. In contrast, Yap proteins activate the expression of genes encoding for cytoskeletal regulators, ECM and focal adhesion components; suggesting a direct role in controlling the morphogenetic behavior of the gastrulating precursors. Quantitative live-imaging analysis of cell displacement trajectories confirmed that Yap proteins are required for dorsal migration towards the midline. By following Yap activity *in vivo* using a Tead sensor (i.e. *4xGTIIC:GFP*), we show that yap-depending transcriptional program is triggered in dorsally migrating precursors rather than in compacted cells at the developing axis. In the absence of Yap function, mutant cells show reduced focal contacts and cortical actin recruitment, and fail to acquire the characteristic flattened morphology of the wild type migratory cells. As reported in numerous studies (Aragona et al., 2013; Davis and Tapon, 2019), we confirm that also in the context of gastrulating cells Yap activation depends on acto-myosin contractility. These observations point to the existence of a Yap-dependent mechano-regulatory loop that ensures the efficient convergence of the precursors to the midline; a mechanism likely conserved in many other homeostatic and developmental processes.

## RESULTS

### Yap paralogs are required for proper axis condensation in medaka

Gastrulation relies on massive collective cell migration in order to place the three germ layers in their correct topological position while directing the formation of the embryo body axis (Keller, 2005). A common theme in collective cell movements is a requirement for an interplay between signaling and mechanic cues to drive cytoskeletal rearrangements needed for oriented cell displacement. Previous work in different vertebrate species, including our work in medaka (Vazquez-Marin et al., 2019), hinted to a potential role for the mechanosensor Yap during axis formation and elongation, thus making this transcriptional regulator and its paralogs perfect candidates to orchestrate the genetic programs controlling stereotypic cell rearrangements. To gain insight into a potential role of Yap family proteins in this process, we focused on the phenotypic consequences of mutating both *yap* paralogs, *yap1* and *yap1b* in gastrulating medaka embryos. When we examined *yap1-/-* medaka embryos at stage 20 (hereinafter referred as ‘single mutants’) we observed that the somite formation was affected and the axis was thicker and shorter (Fig 1A, B). Despite these defects, the primary embryo axis was formed and when we observed these same embryos at later stages (stage 24), the somitogenesis recovered and the anterior-posterior (A-P) axis appeared generally normal, just slightly shorter and thicker compared to the A-P axis of wild-type (WT) embryos (Porazinski et al., 2015; Vazquez-Marin et al., 2019) (Fig S1A-D). The defects in the development of *yap1-/-*;*yap1b-/-* double mutants (hereinafter *yap* double mutants) were remarkably stronger, since they did not condense any posterior axis nor form any somites at stage 20 (Fig 1A, C). In addition, these embryos did not survive to later stages, demonstrating the higher severity of the yap double mutants’ phenotype. To further characterize the phenotype, we performed a DAPI and Phalloidin staining, in order to visualize nuclei and filamentous actin respectively (Fig 1D-F, S1E). Confocal analysis showed how in WT embryos A-P axis condensation becomes apparent and actin network concentrates at the epiboly front, particularly at the closing blastopore (Fig 1D-D’’, S1E and Movie 1). In contrast, in *yap* single mutants, actin staining appeared more diluted at the delayed blastopore margin and a decrease density of cells at the midline was observed (Fig 1E-E’’, S1E and Movie 2). In agreement with our previous findings, *yap* double mutants completely fail to ensemble their posterior part, displaying a significantly reduced D-V accumulation of cells at the midline, and without an apparent definition of the presumptive neural plate and paraxial mesoderm masses (Fig 1F-F’’, Fig S1E and Movie 3).

**Figure 1.**
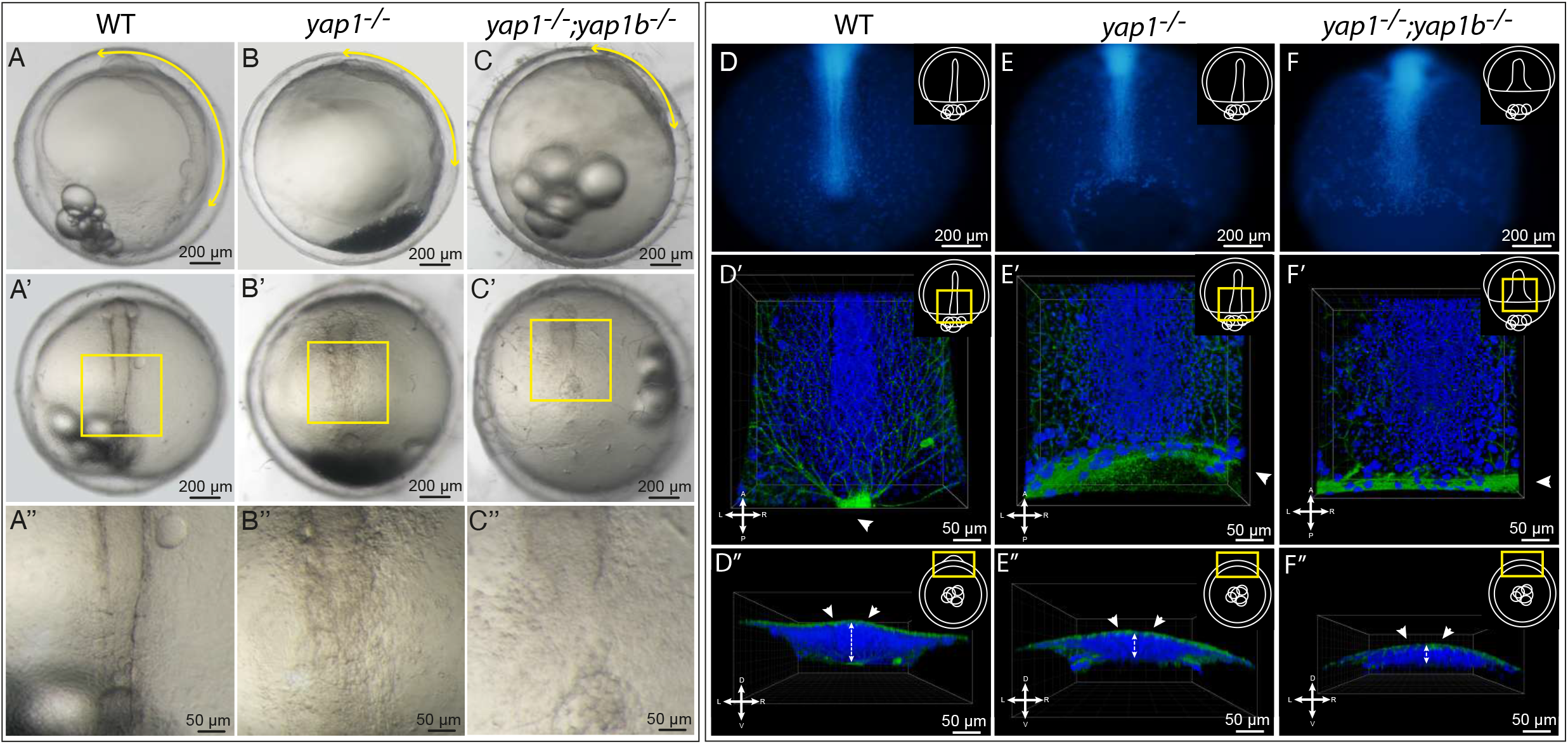
Analysis of *yap* mutants’ phenotype. **(A-C)** Brightfield images of WT, *yap1^-/-^* and *yap1^-/-^;yap1b^-/-^* embryos at stage 20 (post gastrulation stage) (A-C). Yellow double-headed arrows highlight the shortening and the widening of the A-P axis (A-C). A’’-C’’ correspond to the magnifications indicated with a yellow square in their corresponding A’-C’ images. **(D-F)** DAPI staining, in which nuclei are labeled in blue, were performed in WT, *yap1^-/-^* and *yap1^-/-^;yap1b^-/-^* embryos fixed at stage 16-17 (late gastrula) (D, E and F). Whole embryos are shown under the fluorescent stereo microscope. DAPI and Phalloidin immunostained confocal images show the posterior axis of WT, *yap1^-/-^* and *yap1^-/-^;yap1b^-/-^* embryos at stage 16-17 (nuclei in blue and filamentous actin in green) (D’, E’ and F’). XZ projections of the DAPI and Phalloidin immunostaining, showing the width of the A-P axis of WT, *yap1^-/-^* and *yap1^-/-^;yap1b^-/-^* embryos (D’’, E’’ and F’’) Embryos’ orientations are indicated with a cross (A: anterior, L: left, R: right, P: posterior). Yellow rectangles in schematic embryo representations indicate the area depicted in each image. Scales bars 200 μm (A-C, A’-C’ and D-F) and 50 μm (D’-F’ and D’’-F’’).

Yap has a crucial role in cell survival (Wang et al., 2020; Yosefzon et al., 2018), and an increase in cell death in yap single and double mutants has been reported in post-gastrulating embryos at stage 20-22 (Porazinski et al., 2015; Vazquez-Marin et al., 2019). Therefore, we first evaluated if a possible explanation for the observed phenotype was an increase in cell death in *yap* double mutants during gastrulation. To this end, we quantified the number of apoptotic cells labeled by caspase 3 in WT and *yap* double mutants in late gastrulation embryos at stage 16. We did not observe significant differences in cell death density between WT and *yap* double mutant embryos at these earlier stages of development (Fig S1F, G). Thus, an increase in cell death is not behind the observed failure to condense the posterior axis in *yap* double mutants.

The unique phenotype here described indicates that Yap proteins cooperate to control the condensation of the primary embryo axis. Given that cell death could be ruled out as a potential cause for the abnormal cell staking, we decided to explore alternative mechanisms behind the gastrulation and axis condensation defects.

### Yap is needed for a correct cell migration during gastrulation

In teleost embryos, axis condensation is achieved by dorsal migration and lateral intercalation of the precursors at the midline (Solnica-Krezel and Sepich, 2012). Since we observed a clear failure in midline cell stacking in our *yap* mutants, we asked ourselves if directed cell migration was as well altered. To assess that, we analyzed cell trajectories during gastrulation using live imaging in medaka embryos. In WT embryos, cells move dorsally from the lateral regions towards the A-P midline in a very straight and targeted manner (Fig 2A and Movie 4). *yap* single mutants seem to display lower accuracy in their directionality and cells are slightly delayed when reaching the A-P midline (Fig 2B and Movie 5), defects that are largely compensated as development progresses. Displacement defects are markedly accentuated in *yap* double mutant embryos, where many cells display abnormal trajectories and deficient migration towards the midline (Fig 2C and Movie 6). To further confirm these observations, we performed high throughput analysis of cell tracking to measure the main parameters involved in the directed migration of cells. First, we measured cell displacements, which quantifies how much a cell moves from its start point to the midline, and represented this parameter with a color gradient (Fig 2D, E). We could observe that unlike in WT embryos, large-displacing cells (i.e. red and yellow trajectories) were rarely detected in *yap* double mutants, whereas small-displacing cells (i.e. blue and green trajectories) predominate (Fig 2D, E). The statistical analysis of these measurements confirmed our observations (Fig 2F); not only all the mean values of displacement were significantly lower in *yap* double mutants compared to WT embryos (n=3), but also the outliers observed in each WT, which represent cells with large displacement, were not detected in *yap* double mutants. The second parameter we evaluated was the migratory persistence of cells, which measures how long a cell keeps the same directionality. This analysis revealed that the migratory persistence of *yap* mutant cells is significantly reduced compared to WT cells (Fig 2G). Finally, we also measured the cell trajectory length, which quantifies the total distance that a cell moved and their mean velocity. Similarly, to the previous parameters, we could clearly observe that WT cells move faster and longer than *yap* mutant cells (Fig 2H, I). Taken together, these results indicate that Yap proteins play an essential role to direct cell migration in gastrulating embryos.

**Figure 2.**
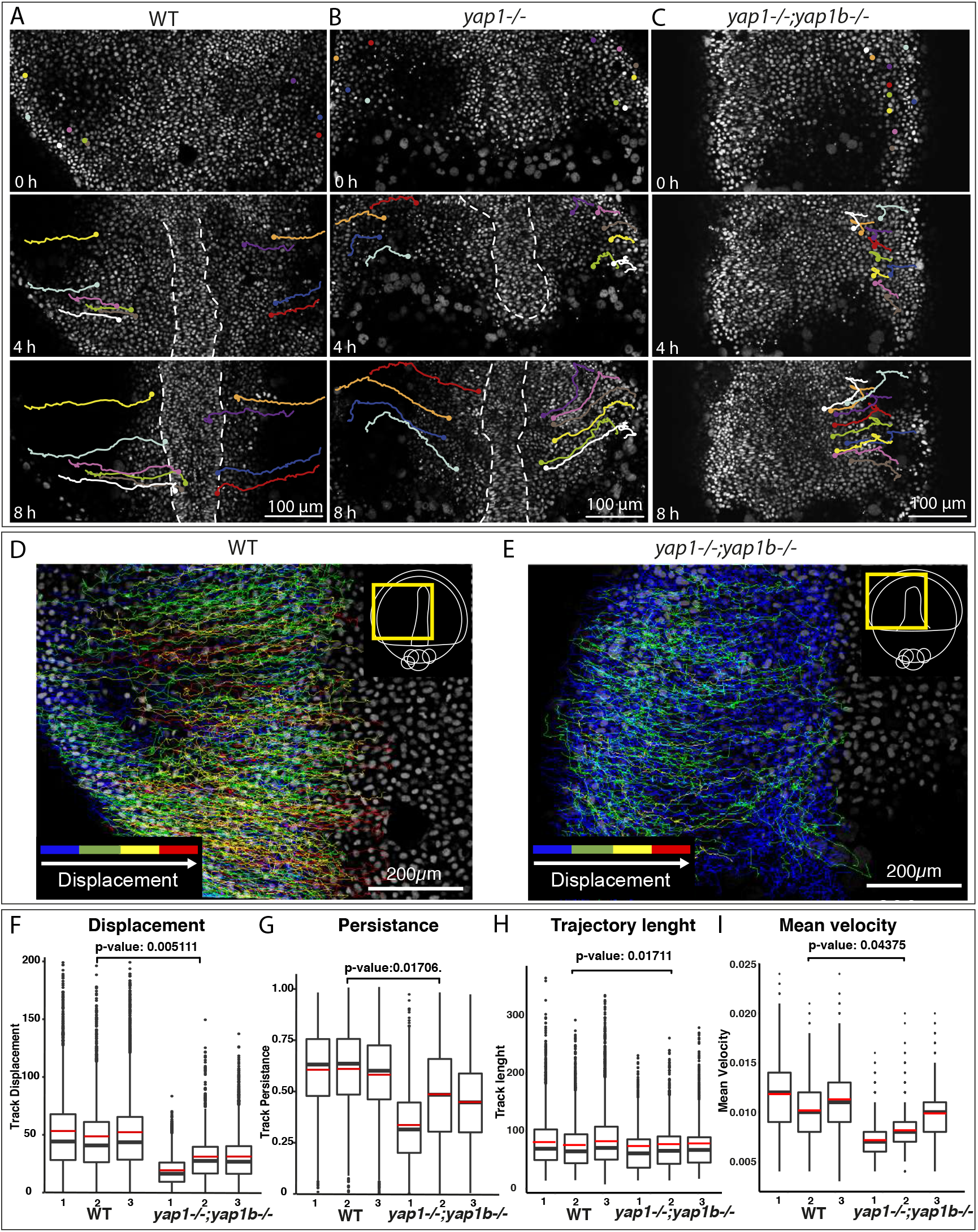
Defective directed cell migration in gastrulating yap mutant embryos. **(A-C)** Manual tracking of representative cells of gastrulating WT (A), *yap1^-/-^* mutant (B) and *yap1^-/-^;yap1b^-/-^* mutant embryos (C) injected with *Histone2B*::GFP for nuclei visualization. They are still images from movies 4-6 at 0h, 4h and 8h. Each cell trajectory is represented with a color line. When possible, the midline is represented white dashed lines. **(D-E)** Total individual cell migratory tracks over 8 h in WT and *yap1^-/-^;yap1b^-/-^* embryos. The color code of the trajectory lines indicates the cells’ displacement (distance between the start and end position of a cell), where blue represents the smallest and red the largest displacement. Yellow rectangles in schematic embryo representations indicate the area depicted in each image. **(F)** Quantification of cell displacement in WT and *yap1^-/-^;yap1b^-/-^* embryos. P-value = 0.005111. **(G)** Quantification of cell migratory persistence, measuring for how long a cell keeps the same direction of movement, in WT and *yap1^-/-^;yap1b^-/-^* embryos. P-value = 0.01706. (G) Quantification of cell trajectory length, measuring the total length of a cell trajectory (H). P-value = 0.01711. (I) Quantification of the cell mean velocity, measuring distance between two cell’s positions divided by the time difference, in WT and *yap1^-/-^;yap1b*^-/-^embryos. P-value = 0.04375. Boxes represent the quartiles; the whiskers indicate the maximum and minimum values. Red and black lines indicate the median and the mean, respectively. To analyze whether experimental groups were significantly different, a variance test followed by a two-sided Student’s t tests were performed on the means of WT and *yap1^-/-^;yap1b^-/-^* embryos. Scale bars 100 μm(A-C) and 200 μm (D-E).

### Yap transcriptional programs primarily regulate cytoskeleton organization and cell adhesion components

Diverse molecular cues have been shown to direct polarized cell movements during gastrulation, such as cell-to-cell adhesion, interaction with the Extracellular Matrix (ECM), or chemotaxis (Solnica-Krezel and Sepich, 2012). On the other hand, Yap proteins have been shown to activate multiple transcriptional programs that are context-dependent (Zanconato et al., 2015). Therefore, in order to get a complete picture on how Yap might be directing cell trajectories, we performed a comparative RNA-seq analysis of WT, *yap* single and double mutant embryos at mid-late gastrulae stage (stage 16) (Fig 3, S2, S3; Table 1). Using this approach, we identified 717 and 1178 differentially expressed genes (DEGs) in *yap* single and double mutants compared to WT embryos, respectively (Fig 3A, S3A; Table 1; Supplementary dataset 1). Principal components analysis (PCA) of the obtained results showed that WT samples cluster together, whereas *yap* single and double mutants group together (Fig S3B), supporting our previous finding that both *yap1* and *yap1b* control very similar transcriptional programs (Vazquez-Marin et al., 2019). For that reason, from now on, we will focus on the analysis of the yap double mutants most severe phenotype.

**Figure 3.**
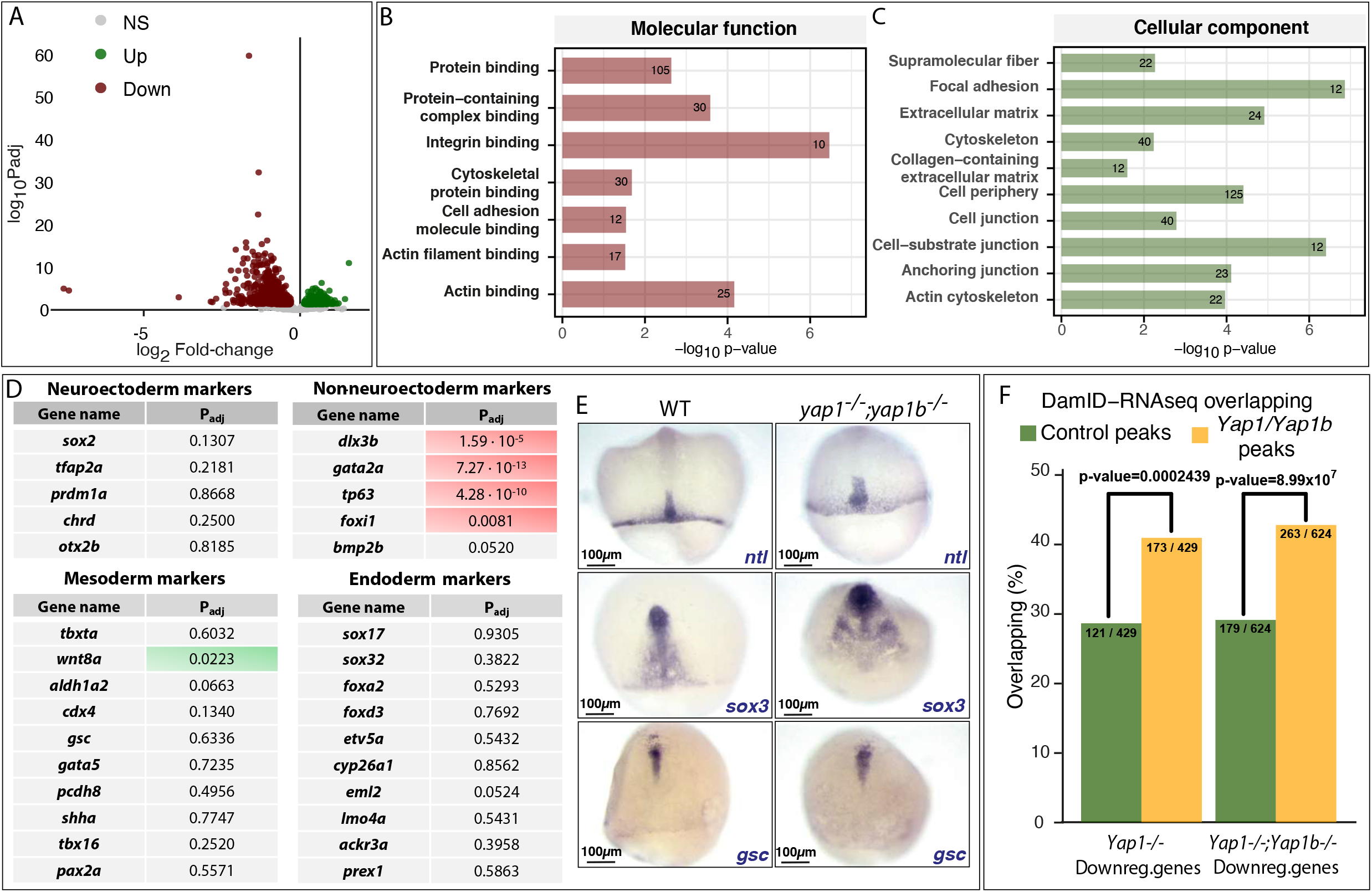
Yap-dependent transcriptional programs. **(A)** Volcano plot graph showing the differentially expressed genes (DEGs) obtained when comparing WT to *yap1^-/-^;yap1b^-/-^* embryos. Gray dots: no differentially expressed genes; Green dots: up-regulated genes in *yap1^-/-^;yap1b^-/-^* embryos compared with WT; Red dots: down-regulated genes in *yap1^-/-^;yap1b^-/-^* embryos compared with WT. Differential gene expression analysis was carried out using the R package DESeq2 (padj < 0.05; -log FC = 1). **(B,C)** Gene Ontology (GO) enrichment of the DEGs in *yap1^-/-^;yap1b^-/-^* embryos compared with WT, classified in molecular function (B) and cellular component (C). **(D)** Differential expression of the 10 most conserved markers of each germ layer in *yap^-/-^;yap1b^-/-^* compared to WT embryos. Adjusted p-values are shown. Red indicates significantly down-expressed genes in *yap1^-/-^;yap1b^-/-^* embryos compared with WT; Green indicates significantly up-expressed genes in *yap1^-/-^;yap1b^-/-^* embryos compared with WT. **(E)** *In situ hybridization* analysis of the expression of *ntl* (mesodermal marker), *sox3* (ectodermal marker) and *gsc* (endodermal marker) in WT and *yap1^-/-^* and *yap1*^-/-^embryos at st 16. Scale bars 100 μm. **(F)** Quantification of the overlap between DamID peaks obtained previously (Vázquez-Marín et al., 2019) and the peaks from the comparative RNA-seq data of WT versus *yap1^-/-^* (P-value = 0.0002439) or *yap1^-/-^;yap1b^-/-^* (P-value = 8.99 × 10^7^) embryos. The statistical significance for the comparison between control and identified overlapping DamID peaks was calculated applying a two-proportion Z-test.

To understand the mechanisms behind Yap activity, we next studied Gene Ontology (GO) terms enrichment in the DEGs in *yap* double mutants *vs* WT (Fig 3B, C and S2C, D). We explored four different GO categories: molecular function, cellular component, biological process and KEGG pathway. In the molecular function category, we identified integrin binding as the most significantly enriched GO term, followed by others such as acting binding or cell adhesion (Fig 3B). Very consistent results were also obtained for the remaining GO categories (Fig 3C and FigS2C, D).

Thus, significantly enriched GOs terms identified in *yap* double mutants were related to cell-ECM adhesion (i.e. focal adhesion (FA) or collagen-contain ECM), Hippo signaling, and actin cytoskeleton regulation and organization) (Fig 3C, Fig S3C, D; Supplementary data set 2). Similar GO enrichments were obtained when DEGs between yap single mutants *vs* WT were considered (Fig S3E-H). These data indicate that Yap paralogs primarily regulate the expression of actomyosin cytoskeleton and ECM-cell adhesion components (Supplementary dataset 2).

Cellular rearrangements during gastrulation are often coordinated with lineage restriction and germ layers specification. To verify whether cell-fate is compromised in *yap* mutants we examined in more detail our RNA-seq datasets, which in principle did not yield significantly enriched GO terms consistent with that hypothesis (Fig3 B-C, Fig S3 C-H). For further confirmation, we checked the expression of a battery of 10 conserved specifiers of each germ layer. No significant differences were observed for key mesoderm and endoderm markers when their expression was compared between yap mutant and WT embryos (Fig 3D). Similarly, no significant differential expression could be detected when neuroectoderm markers were examined. The only genes that appeared significantly downregulated correspond to non-neural ectoderm specifiers, a cell identity that is acquired once gastrulation has ended (Fig 3D). In order to confirm these observations, we compared the expression patterns of a mesoendodermal marker, *no-tail* (*ntl*), an endodermal marker, *goosecoid* (*gsc*), and a ectodermal marker, *sox3* (Fig 3E). In agreement with our RNA-seq data, these three germ-layer markers do not appear down-regulated in *yap* mutants when compared to WT embryos. However, their expression patterns appeared wider in *yap* mutants, which suggests a failure in cell condensation, in line with the defective migration of cells that we observed (Fig 3E). Taken together, these analyses point to a failure in the activation of the genetic program controlling cytoskeleton reorganization and cell adhesion, rather than a problem in cell-fate acquisition, what is behind the cell migration defects.

To gain further insight into the genetic program controlled by Yap proteins, we compared our RNA-seq data with DamID-seq results we previously obtained in stage 16 medaka embryos (Vazquez-Marin et al., 2019). Using the DamID-seq technique, we had generated maps of chromatin occupancy for Yap1 and Yap1b in gastrulating embryos. Then, by cross-comparing genes neighboring Yap paralogs binding sites with our list of DEGs, we could determine which of these genes are potentially direct targets of Yap. We observed that a significantly high percentage of these DEGs are direct targets of Yap1/Yap1b (Fig 3F). Importantly, among the direct targets, we could confirm relevant regulators of the cytoskeleton, such as *marcksl1b*; structural ECM encoding genes, such as *lamc1*; well-known Yap targets *cccn1/cyr61*; and Yap regulators such as *src* (Fig S3 and Supplementary dataset 2).

### Yap is active in migratory cells converging to the midline

For a cell to migrate it needs to polymerize actin at the leading edge to drive protrusions that adhere to the substrate through focal adhesions (FAs) (Shellard and Mayor, 2020). Our results suggested that Yap activates the transcriptional programs controlling cytoskeleton and focal adhesion components required for proper cell migration during gastrulation. Thus, we wanted to check the spatiotemporal dynamics of Yap activation by following both the transcriptional activity of a Tead sensor, and the expression of *marcksl1b* a bona-fide Yap target as determined by DamID-seq and RNA-seq analyses (Vazquez-Marin et al., 2019).

Tead is a family of transcriptional regulators that interact with Yap proteins acting as main mediators of the Yap-dependent response (Stein et al., 2015; Vazquez-Marin et al., 2019; Zanconato et al., 2015; Zhao et al., 2008). To monitor Tead expression, and therefore the activation of Yap, we generated a new medaka transgenic line in which the Tead-responsive *4xGTIIC* enhancer, previously tested in zebrafish, was coupled to GFP (*4xGTIIC::GFP*) (Mahoney et al., 2005; Miesfeld et al., 2015). To validate this transgenic line, we first checked that the GFP signal was absent in yap mutants (Fig S4A). Then, by injecting Yap mRNA fused to the mCherry reporter gene (Yap::mCherry), we confirmed that the cells with higher levels of nuclear Yap::mCherry are the ones displaying the higher intensity of the Tead reporter signal (Fig S4B). These results corroborate that the *4xGTIIC::GFP* transgenic line respond specifically to Yap signaling. Using this tool, we could then follow in vivo the dynamics of Yap activation during gastrulation by live confocal imaging (Movie 7). Interestingly, we observed that Yap is active in the cells that are actively migrating towards the midline, rather than in the midline itself, where cell density is higher (Movie 7 and Fig 4A). To corroborate these observations, we also examined the expression pattern of *marcksl1b* by *in situ* hybridization. Marcksl1b is annotated as a protein involved in promoting cell motility by regulating actin cytoskeleton dynamics as well as filopodium and lamellipodium formation (El Amri et al., 2018). Matching our observations with the Tead reporter line, we saw that in gastrulating embryos, *marcksl1b* is expressed mainly in the lateral cells actively converging towards the midline (Fig 4B). Once gastrulation is completed, we also observed a new pattern of expression, with *marcksl1b*-expressing cells proximal to the midline (Fig 4B). As expected, *marcksl1b* expression is absent in *yap1* mutant embryos (Fig S4C). These two sets of results stress the fact that during gastrulation, Yap is specifically active in cells that are moving towards the midline, while inactive in the more static and compact cells at the midline (Fig4C, D). Thus, we concluded that the migration defects we observed in gastrulating *yap* mutant embryos must be caused by the absence of Yap activation in the migratory cells.

**Figure 4.**
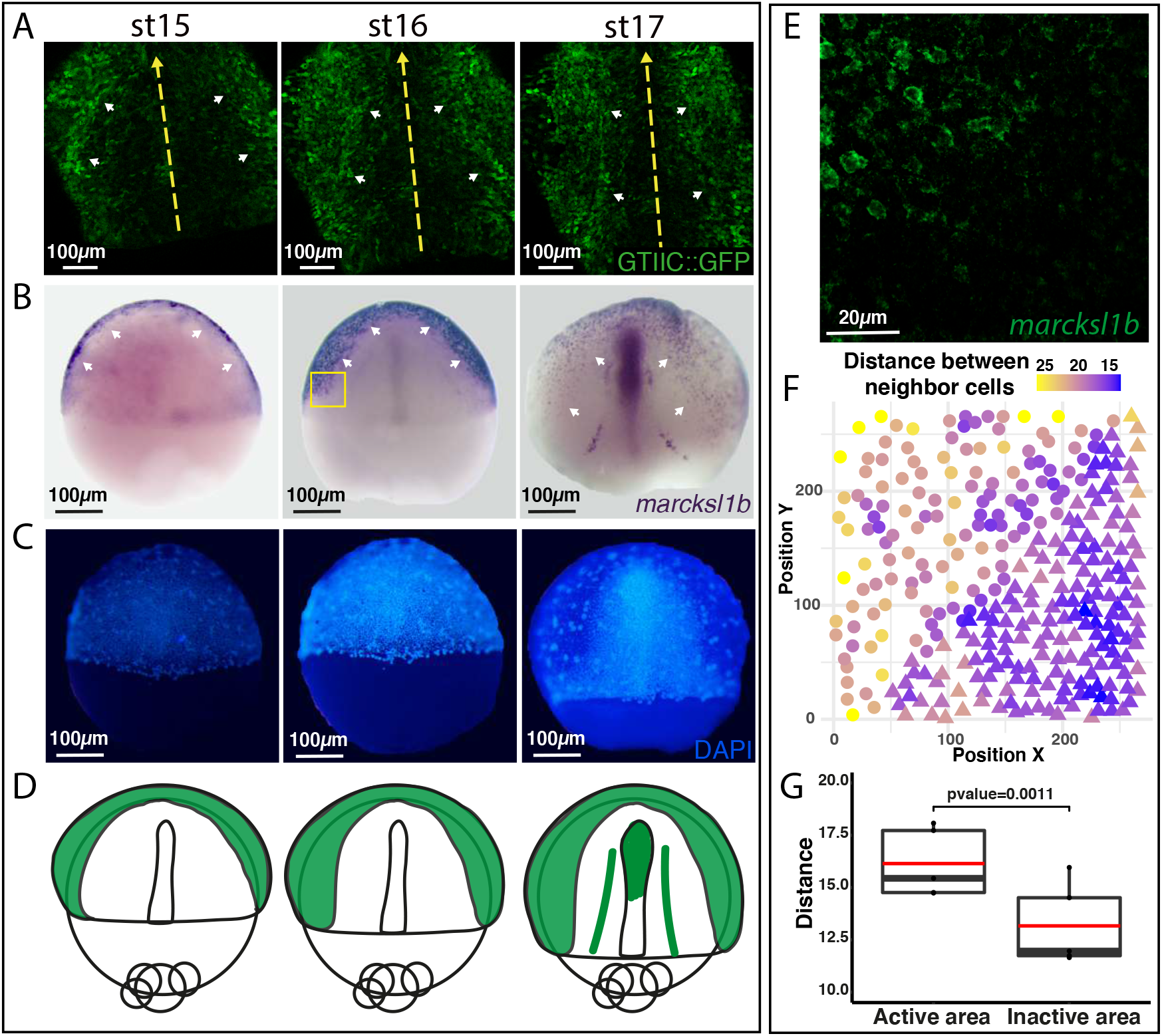
Yap activity dynamics during gastrulation. **(A)** *GTIIC::GFP* transgenic embryos are shown under fluorescent confocal microscope at stages 15, 16 and 17. White arrowheads point to the GFP-positive cells Yellow arrows indicate A-P. They are still images from movie 7. **(B)** *In situ hybridization* analysis of the expression of *marcksl1b* in WT embryos at 15, 16 and 17. White arrowheads point to the cells expressing *marcksl1b*. **(C)** DAPI staining of the embryos in (C). **(D)** Schematic representation of the expression of *marcksl1b* (Yap activation) in green. **(E-F)** Fluorescent confocal microscope image of *marcksl1b* fluorescent *in situ hybridization* of the zone were *marcksl1b* expression decreases in WT embryos at st 16 (E) and its corresponding cell density analysis (F). The region depicted in (E) is equivalent to the region marked with a yellow rectangle in the st 16 embryo in (B). The XY position of the nuclei’s centroids were represented. Circle: nuclei localized in a *marcksl1b* positive area; Triangle nuclei localized in a *marcksl1b* negative area. The color gradient refers to the mean of the distance between a cell and its five closest nuclei. **(G)** Quantification of the mean distance between nuclei in the *marcksl1b*-positive area (Active) and in the *marcksl1b*-negative area (Inactive). P-value = 0.0011. Boxes represent the quartiles; the whiskers indicate the maximum and minimum values. two-sided Student’s t-test was used to evaluate statistical significance. Scale bars 100 μm (A-C) and 20 μm (E).

In agreement with our previous findings (Vazquez-Marin et al., 2019), we confirmed that *yap1* is ubiquitously expressed at early gastrula (stage 15), and as gastrulation advances, the expression is enriched at the condensed midline (stage 16), decreasing in the lateral regions when gastrulation completes (Stages 17 and 18) (Fig S4D). This discrepancy between *yap* expression and Yap activation, likely funded on the highly complex post-transcriptional regulation of the Yap protein (Pocaterra et al., 2020) suggests that Yap signaling inhibition is not transcriptional, but that it rather depends on protein regulation. It has been described in *in vitro* cell assays that Yap activity can be directly inhibited by cell density (Varelas et al., 2010; Zhao et al., 2007). To examine whether this anti-correlation between Yap activation and cell compaction was also significant in the gastrulation context, we examined density maps in relation to *marcksl1b* expression (Fig 4E-G). We could determine that the distance between neighbors is significantly higher (lower cell density) in Yap active-areas than in Yap-inactive areas (Fig 4G), pointing to cell density as a potential modulator of Yap activity during gastrulation.

### Yap impairs cortical actin recruitment and focal adhesions assembly in migratory cells

We previously showed that Yap proteins are active in convergent dorsal cells, in which they modulate the expression of cytoskeletal and ECM-adhesion components. We therefore asked how Yap impinges in the actin cytoskeleton of cells and their adhesion to the ECM to facilitate directed migration. To this end, embryos were injected at one cell stage either with *Utrophin:GFP* mRNA, to label filamentous actin, or *Pax::mKate* mRNA, to reveal FAs assembly. Then the distribution of these tracers was examined using high-resolution microscopy in WT and *yap* double mutants, focusing our attention on the dorsal cells of the inner mass (Fig 5), which have Yap-active and are the ones migrating towards the midline (Fig 4 and S4). Dorsal migrating cells of the inner mass of the embryo display a monolayer distribution sandwiched between the cells of the enveloping layer (EVL), which display a polygonal shape and present remarkably bigger nuclei, and the yolk syncytial layer (YSL). WT inner mass cells present strong accumulation of filamentous actin (in green in Fig5A) and display the spreading shape typical of migratory cells, with extended plasma membrane and protruding filopodia and lamellipodia. In contrast, *yap* mutant cells accumulate less cortical actin and display a rounded shape with less noticeable protrusions (Fig5). We also observed that WT inner mass cells display focal adhesions stripes and foci (in red in Fig5A), which are very reduced in *yap* double mutant cells (Fig5A). This result suggests that Yap is essential for FAs maturation, as these structures enlarge and lengthen as they mature (Pasapera et al., 2010). By looking at the XZ projections of *Paxillin:mKate* in WT cells, we observed that FAs clusters tend to accumulate at the surface contacting the YSL, thus suggesting that inner mass cells are migrating preferential over the yolk surface (Fig5A’). *Yap* mutant cells clearly lack these polarized adhesion clusters (Fig5A’).

**Fig 5.**
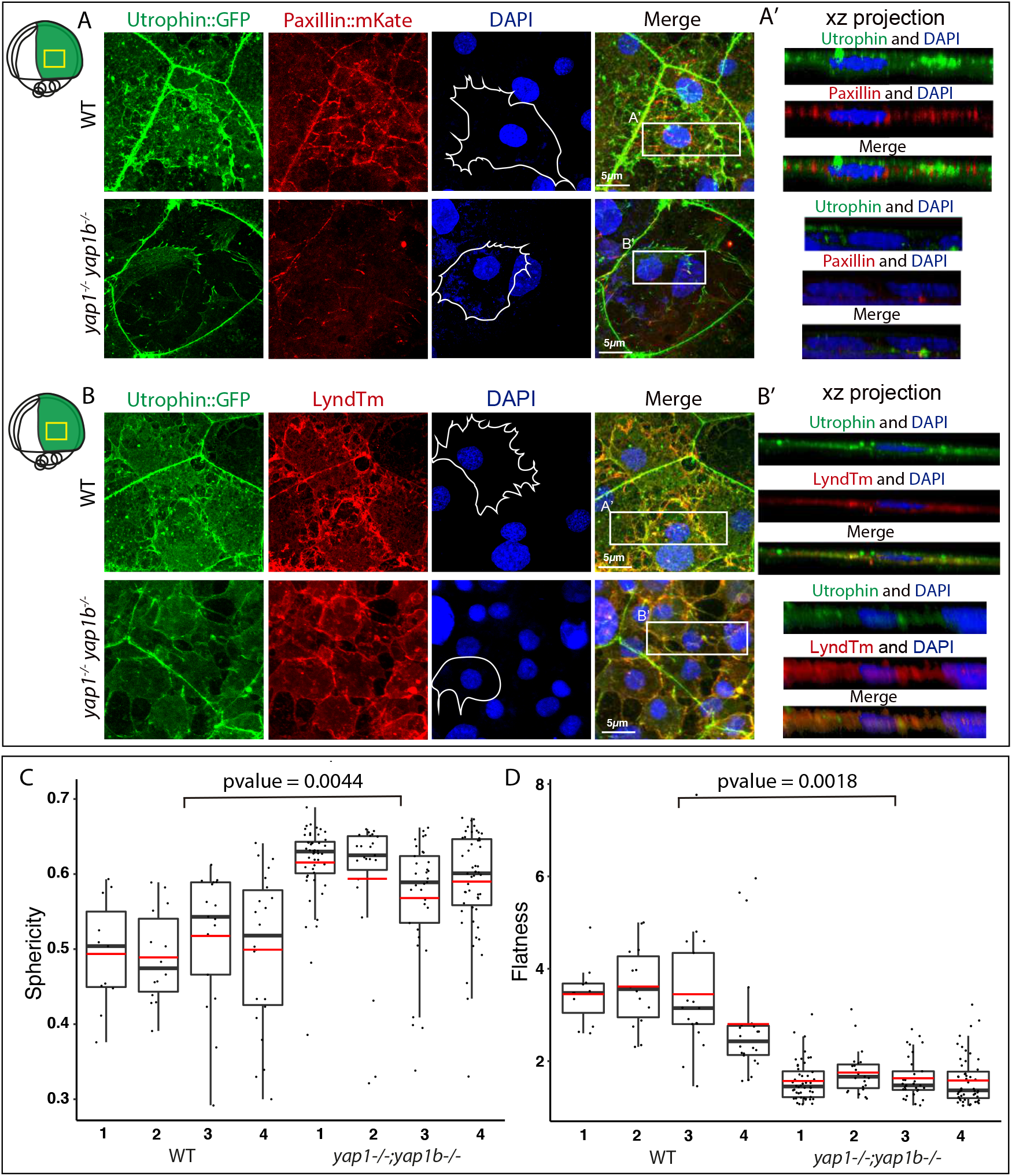
Cell Actin cytoskeleton and adhesions are affected in yap mutants. **(A,B)** Confocal microscopy images of dorsal cells from WT and *yap1^-/-^;yap1b^-/-^* embryos injected with *Utrophin::GFP, Paxillin::mKate* (A), *LyndTm* (B) and stained with DAPI. XZ projections from the sections indicated with white rectangles are shown (A’, B’). Schematic representation of the embryo indicating the area of interest with a yellow rectangle is shown in the upper left side of the panel. Cell shapes are represented with white lines in the images corresponding to DAPI. **(C)** Quantification of dorsal cells’ nuclei sphericity, which measures the degree to which a nucleus approaches the shape of a sphere, in WT and *yap1^-/-^;yap1b^-/-^* embryos. P-value: 0.0004445. **(D)** Quantification of dorsal cells’ nuclei flatness, which refers to the ratio between the second and the third axis of an ellipsoid, in WT and *yap1^-/-^;yap1b^-/-^* embryos. P-value: 0.001808. Boxes represent the quartiles; the whiskers indicate the maximum and minimum values. Two-sided Student’s t-tests were performed to evaluate statistical significance. Scale bars 5 μm.

To get further insight into cells’ morphology, we performed a similar experiment but now injecting *Utrophin:GFP* together with *Lyn-tdTomato (LynTm)* mRNA, to visualize the plasma membrane. These results confirmed the cortical accumulation of filamentous actin (in green in Fig5B) and the expanded protrusions of WT cells, which presented multiple filopodia and membrane ruffles along all the cell surface (in red in Fig5B). In contrast, *yap* mutant cells displayed a more rounded and uniform morphology with remarkably lower number of filopodia.

By examining xz projections of the nuclei (Fig 5B’), we observed that WT nuclei and cells tend to appear flatter than *yap* mutant cells. Since mechanical coupling to the ECM and nuclear deformation are required for Yap nuclear shuttling (Elosegui-Artola et al., 2017; Elosegui-Artola et al., 2016), we decided to explore this observation further. To this end, the 3D nuclei morphology of WT and *yap* mutant cells was measured. We found that two main morphological parameters, sphericity and flatness, were significantly different between WT and *yap* mutant cells; *yap* mutant nuclei are significantly more spherical than those of WT cells (Fig 5C) and display a lower flatness index (Fig 5D). From these measurements, we hypothesized that the noticiable reduction of cortical filamentous actin and FAs observed in *yap* mutant cells may lead to a decrease in intracellular tension, which is reflected in a more relaxed and rounded cell morphology.

All these data suggest that Yap activity is promoting the formation and maturation of FAs and the polymerization of cortical actin in migratory cells. Thus, in the absence of Yap activity cells would be unable to establish mature FAs and actin bundles, essential to respond to ECM cues.

### Yap senses and activates intracellular tension within a positive feedback loop

Our prior results indicated that Yap activity regulates FAs and actomyosin cytoskeleton, which allows coupling intracellular tension to the ECM and facilitates cell spreading. Yap transcriptional regulators have been characterized as a mechanosensor (Aragona et al., 2013; Dupont et al., 2011). Therefore, we wondered if yap is not only required to maintain and transmit intracellular tension, but is also able to sense it, responding to it by promoting its own activation through a positive feed-back loop mechanism. The integrins, as components of the focal adhesions, transmit information about the different rigidity of the ECM to the cells through the interaction with the Rho/Rock pathway, which modify the F-actin cytoskeleton and mechanically activates YAP/TAZ (Elosegui-Artola et al., 2016; Sero and Bakal, 2017; Totaro et al., 2018). To test our feed-back loop hypothesis, we applied the pharmacological inhibitor ROCKOUT, which interferes the mechanosensing Rho/ROCK pathway. When this pathway is inhibited, there is a reduction of stress fiber formation and FA maturation, which translates in reduced intracellular tension (Nardone et al., 2017; Piccolo et al., 2014). We therefore treated gastrulating embryos with ROCKOUT for a short developmental window (2h) and examined *marcks1b* expression, a direct target of Yap according to our data (Fig 3, 4), as a readout of Yap activation. Interestingly, *marcks1b* expression appeared very reduced in ROCKOUT-treated embryos when compared to wild type, thus indicating a diminished transcriptional activation by Yap (Fig 6A). These ROCKOUT-treated embryos also showed a clear delay in epiboly, defect also observed in *yap* double mutants (Fig 6A). Therefore, the pharmacological inhibition of the intracellular tension during gastrulation leads to the inhibition of Yap activity, mimicking *yap* mutants’ phenotype. This result suggests that a direct feedback loop between intracellular tension and Yap activation operates to maintain directed cell migration in gastrulating cells.

**Fig 6.**
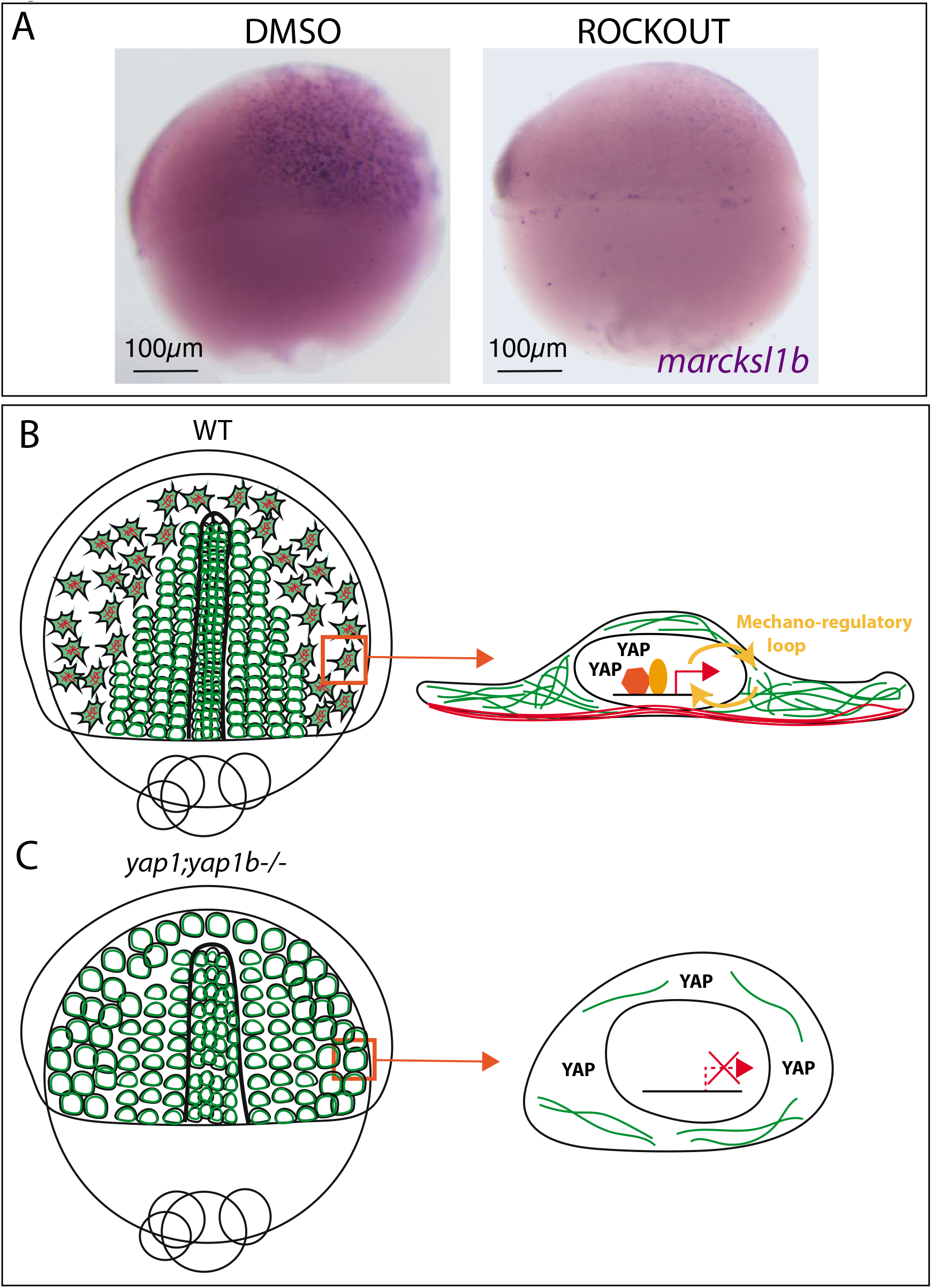
Yap senses intracellular tension within a positive feedback loop. **(A)** *In situ hybridization* analysis distribution of *marcksl1b* in WT embryos treated with DMSO or ROCKOUT drug during 2 h. Lateral images are shown. Scale bars 100 μm. **(B,C)** Schematic representation of the differences between lateral migrating cells converging to the midline in WT (B) and and *yap1^-/-^;yap1b^-/-^* embryos (C).

## DISCUSSION

### Yap is specifically active in dorsal cells migrating towards the midline

Cells need to know where and how to move, not only during embryogenesis, but also for the proper homeostasis and functioning of adult tissues. Identifying the underlying cues of cell migration is key, both for understanding self-organization principles as well as for the identification of molecular targets to fight malignant cell migration. Here, we uncover the role of YAP as a transcriptional hub activating the genetic programs required for directed cell migration during gastrulation.

The formation of the embryo axis in teleosts entails cells gathering at the midline through different morphogenetic processes that vary depending on their position. While ventral and anterior cells, closer to the embryo midline, converge following mainly radial and mediolateral intercalation movements, dorsal cells away from the midline converge via directed cell migration (Solnica-Krezel and Sepich, 2012; Wallingford et al., 2002). We showed that YAP is active just in dorsal cells that are far from the midline and need to undergo long distance migrations. Thus, we hypothesize that YAP will be in charge of activating the transcriptional programs required to modify the actomyosin network and cell adhesion required for long-distance displacement. In contrast, at higher cell densities, cells closer to the midline would move mostly by mediolateral intercalation in a YAP-independent manner, and relying instead on Wnt-PCP signaling among others (Heisenberg et al., 2000; Jessen et al., 2002; Wallingford et al., 2000).

We observed a significant correlation between YAP activity silencing and increased cell density closer to the embryo midline. Therefore, we propose that the shape changes imposed by the spatial restrictions in more crowded areas (e.g. cells reduce their size and become rounded, with less contact protrusions), might facilitate Hippo-mediated YAP inhibition. This hypothesis is supported by diverse observations where YAP/TAZ get phosphorylated by Hippo pathway kinases in response to an increase in cell density, getting retained at the cytoplasm and preventing their interaction with their nuclear partners (Aragona et al., 2013; Zhao et al., 2007). Aragona et al., reported that the main determinant for YAP/TAZ inhibition is actually to accommodate to a smaller cell size. Small cells attach to a smaller ECM substrate area, displaying “decreased integrin-mediated focal adhesions, reduced actin stress fibers, and blunted cell contractility” (Aragona et al., 2013). Importantly, these mechanical cues have been shown to be essential for YAP/TAZ nuclear localization and activity (Dupont et al., 2011; Wada et al., 2011). Cells also become rounded in other scenarios, such as culturing cells in soft substrates, placing them in suspension, or disrupting the F-actin cytoskeleton. Interestingly, in all this cases, YAP/TAZ signaling is inhibited due to a decrease in the mechanical constrains of the cytoskeleton, independently of changes in cell-to-cell contacts (Aragona et al., 2013; Grannas et al., 2015; Wada et al., 2011; Zhao et al., 2012). The work of Elosegui-Arteaga et al. illustrates how cytoskeletal and cell shape changes affect YAP activation in the opposite cell configuration, i.e. spreading cells. They describe how cell flattening triggers nuclear pores relaxation, allowing transcription factors, such as YAP, to enter the nucleus upon cell deformation (Elosegui-Artola et al., 2016). Our results show that YAP-activated cells display the typical spreading shape, with more flattened and less spherical nuclei. In line with that, we saw that the YAP mutant cells are less flatten and more spherical.

### Yap1 paralogs direct the changes in cell cytoskeleton and ECM adhesions that allow their proper migration during gastrulation

According to our observations YAP plays a purely morphogenetic role at gastrulation stages. We did not see, however, an involvement of YAP in other developmental processes such as cell proliferation and cell death, as it has been shown in fish at later stages of development (Porazinski et al., 2015; Vazquez-Marin et al., 2019). A very recent work shows that the ancestral role of YAP in metazoans is precisely in cytoskeletal dynamics rather than in cell proliferation (Phillips et al., 2021), suggesting that YAP has independent roles depending on the specific temporal and spatial context of the cell. This is also in line with different observations in cell culture, which identified different biological processes related to YAP depending on the cell type (Nardone et al., 2017; Zanconato et al., 2015). Our observations are in concordance with one of these works, in which the authors describe that when cells lack YAP activity, they lose their ability to adhere to the ECM, they present a decreased number of FA spikes and they cannot spread over the ECM (Nardone et al., 2017).

The role of Yap and its paralogs during morphogenesis has been previously studied in teleost models. However, their role in cell migration at earlier gastrulation stages has never been analyzed before. A first work in medaka showed that single *yap1* mutants present pronounced body flattening, delay in blastopore closure and misalignment defects (Porazinski et al., 2015). In accordance with our results, the authors suggest that the morphogenetic phenotype is caused by a reduction of the actomyosin-mediated tension. They did not observe, however, any defect in gastrulation movements and axis condensation, as the *yap1b* paralog was still functional. This is in line with our previous observation that both paralogs, *yap1* and *yap1b*, are functionally redundant and that *yap1b* can partially rescue *yap1* suppression early during development (Vazquez-Marin et al., 2019). A second study analyzed the role of *yap1* and its paralog *wwtr1* (also known as *taz*) in zebrafish embryos. In this work, double mutant embryos show a defect in elongating the posterior part of the embryo, by regulating the deposition of Fibronectin (a main ECM component) in the presumptive epidermis (Kimelman et al., 2017). The main difference with our results is that the phenotype observed in zebrafish was first evident at 15-16 somite stage (i.e. much after gastrulation is completed) and did not interfere with the assembly of the primary embryo axis or the somites’ condensation. Given that mild axial defects have also been observed in *yap/wwtr1* double mutants, it will be premature to rule out a role for Yap proteins during gastrulation in zebrafish. A logic assumption is that Yap signaling cooperates with other mechanisms to direct cell migration during gastrulation, and thus it is possible that its role has remained elusive in zebrafish due to compensatory mechanisms. Additionally, it is important to take into account that the spatial configuration of the gastrulating cells varies between medaka and zebrafish embryos, due to the much larger size of the yolk in medaka. Dorsal cells in medaka embryos are flatter and have to travel larger distances to arrive to the midline compared to their equivalent in zebrafish. Therefore, it is possible that the particular geometry of the medaka gastrula has allowed us to uncover the role of Yap proteins in directed cell migration during gastrulation.

### Yap transcriptional program during gastrulation control the expression of genes encoding for cytoskeletal regulators, ECM and focal adhesion components

Our bulk RNA-seq results suggest a tight relationship between Yap activation and an increase in the expression levels of genes mostly involved in cell adhesion, cytoskeleton organization and cell migration. Furthermore, a significant proportion of these identified genes are direct targets of *yap1*/yap1b, according to our previous DamID-seq datasets (Vazquez-Marin et al., 2019). This suggests a straightforward regulatory role in cell adhesion and cell migration during gastrulation. Among genes activated by Yap, we can identify a number of them with a previously identified role in gastrulation, particularly in controlling convergent-extension movements in zebrafish. This is the case of genes encoding for integrins (Ablooglu et al., 2010) and ECM molecules (Latimer and Jessen, 2010), as well as cytoskeletal regulators such as members of the *marcks* family (Iioka et al., 2004), *akap12b* (Weiser et al., 2007), *rock2b* (Marlow et al., 2002) or *vangl2* (Roszko et al., 2015). Besides these gastrulation-related genes, the transcriptional program controlled by Yap during comprises a list of common ‘beacon’ genes, which have been proved to be direct transcriptional targets of Yap/Tead complexes in previous studies, regardless of the cell type and developmental stage considered. This list includes genes such as *ctgfa, cyr61, amotl2b, lats2*, or *col1a1b* (Lin et al., 2017; Vazquez-Marin et al., 2019; Zanconato et al., 2015). In agreement with this, despite the very different cellular context, many of the genes we identified as downregulated in *yap1/yap1b* medaka mutants were also found as differentially expressed in *yap/wwtr1* zebrafish mutants (Kimelman et al., 2017) (e.g. *amotl2b, cyr61, cdc42ep, sorbs3, ctgfa, col1a1b* and *pcdh7*).

### YAP senses and generates mechanical tension in a mechano-regulatory loop

The way that, within tissues, forces are generated, sensed and transmitted is increasingly being understood as a continuous interplay between the cells and their environment. This results in regulatory feedback loops in which cells perceive mechanical cues and respond in turn modifying its own mechanical properties (Hannezo and Heisenberg, 2019; Petridou et al., 2017). In the case of Yap, the general agreement is that the cytoskeletal organization reflects the mechanical state of the tissue and serves as a universal Yap activator; while Yap will transform these inputs into transcriptional changes inducig more cytoskeletal rearrangements (reviewed in (Totaro et al., 2018). Our results suggest that, in the context of gastrulation, Yap is engaged in a positive feed-back loop. We show that Yap activation depends on Rho/Rock-generated cell tension, and in turn triggers a genetic program that by maintaining the spreading shape of gastrulating cells will contribute to sustain intracellular tension levels.

Yap-dependent feedback mechanisms have been described in diverse cellular contexts, ranging from cardiomyocyte regeneration (Morikawa et al., 2015), lung branching in mice (Lin et al., 2017), endothelial cells migration (Mason et al., 2019), and mesenchymal stem cell cultures (Nardone et al., 2017). What our results and all these examples have in common is that mechanically activated Yap/Taz promotes F-actin, FA expression and assembly, and integrin expression; all essential components of the ECM-cell communication (reviewed in (Totaro et al., 2018). These feedback loops confer robustness to the morphogenetic processes and integrate them into their organismal context. Here we have identified YAP as a transcriptional hub that orchestrate the cytoskeletal changes required for a cell to move long distances and arrive on time to their final destination during gastrulation.

## METHODS

### 3D reconstruction of *yap1/yap1b* mutant embryos

Wild type, *yap1* single mutant and *yap1/yap1b* double mutant siblings at stage 17 were fixed with PFA 4% at 4ºC for 2-3 days. Samples were washed extensively in PBS-0.2% Tween and stained with phalloidin Alexa-488 (Invitrogen) in PBS-0.2% Tween solution supplemented with 5% DMSO (1:50) o/n at 4ºC. After extensive washing steps with PBS-0.1% Tween, samples were stained with DAPI (1:1000), mounted in FluoroDish 35 mm plates (WPI) and imaged in a Leica SP5 microscope using a 20x multi-immersion objective. Embryos were imaged dorsoventrally taking images of 50 stacks of 3 μm-length each with a pixel size of 0.379 x 0.379 μm^2^. Tridimensional models were acquired using Imaris 8. After imaging the embryos, each one of them was individually genotyped.

### Whole-mount embryo immunostaining

Embryos collected from *yap1^+/−^;yap1b^+/−^* adult fishes were fixed at stage 16 using 4% paraformaldehyde (PFA). Fixed embryos were dechorionated with forceps. Embryos were washed with P PBS-0.2% Tween, treated with cold acetone at −20 °C for 20 min, then incubated with freshly prepared blocking solution (2% normal goat serum and 2 mg/mL bovine serum albumin (BSA) in PBS-0.2% Tween at room temperature (RT) for 2 h. A purified primary rabbit anti-active caspase-3 antibody (BD Biosciences, 559565) was diluted 1:500 in blocking solution and embryos were incubated overnight at 4 °C. Embryos were then subsequently washed with PBS-0.2% Tween and incubated overnight at 4 °C in the dark with and the Alexa Fluor TM 555 Goat anti-rabbit antibody (Invitrogen #A32727), diluted 1:500. Finally, embryos were washed with PBS-0.2% Tween and incubated overnight at 4 °C with DAPI (Sigma) diluted 1:1000 in PBS-0.2% Tween. For imaging, embryos were embedded in 1% low-melting-point agarose and mounted in FluoroDish 35 mm plates. Confocal laser scanning microscopy was performed using an LSM 880 microscope (Zeiss). Images were processed using ImageJ (Schindelin et al., 2012). For quantification of apoptotic cells, masks were applied for both channels. The mask generated for the red channel was segmented using the Watershed algorithm. Only the regions marked with the primary antibody that also corresponded to nuclei with a 6-200 μm2 area were considered to avoid debris. Apoptotic cells were counted on the embryo surface and extrapolated to a section of 1 mm2. After imaging, embryos were genotyped by PCR to identify yap1/yap1b-related genotypes. To analyze whether experimental groups were significantly different, two-sided Student’s t-tests were performed.

### Analysis of cell movements during gastrulation

Wild type, *yap1* single mutant and *yap1/yap1b* double mutant siblings were injected at onecell stage with H2B-GFP mRNA (Addgene, #53744) at a final concentration of 25 ng/μL. The embryos were incubated for 3 h at 28ºC and o/n at 25ºC. The most promising candidates were then selected the day after in a fluorescent binocular and dechorionated following a three-step protocol with minor modifications (Porazinski et al., 2010). First, embryos were rolled in sandpaper (2000 grit size, waterproof) to weaken their outer structure. Then, the embryos were incubated for 30 min at 28ºC in pronase at 20 mg/mL. Finally, after several washing steps, embryos were incubated for 60 min in hatching enzyme at 28ºC and transferred into a Petri dish with BSS 1x medium supplemented with penicillin-streptomycin and 1-heptanol 3.5 mM (Sigma) to block contractile rhythmical movements. Overnight movies (8-9 h) were acquired using a Leica SP5 microscope. Frames with a pixel size of 0.189 x 0.189 μm^2^ were taken every 4 minutes. Manual cell tracking analysis was carried out using ImageJ (Schindelin et al., 2012). Alternatively, a more precise, semi-automatic cell tracking was also performed using TrackMate (Tinevez et al., 2017). The resulting data were analyzed using R. Cells tracked in less than 15 frames and/or localized initially near the midline were excluded. Two-sided Student’s t-tests were performed to estimate the statistical significance among the different experimental conditions.

### RNA-seq

#### Library preparation

Individual wild type, *yap1* single mutant and *yap1/yap1b* double mutant embryos at stage 16 were homogenized in TRIzol (Ambion). Samples were centrifuged at full speed and the supernatant was transferred to a fresh tube. Chloroform was then added to split the RNA (upper aqueous phase) from the DNA fraction (lower phase). The DNA fraction was precipitated adding glycogen and 100% ethanol and incubating it at RT for 20 min. This fraction was then centrifuged at maximum speed for 30 min at 4ºC. After three washing steps using 75% ethanol, the DNA pellet was then resuspended in 30 μL of TE buffer. The RNA fraction was precipitated adding RNA-grade glycogen (ThermoFisher Scientific) and isopropanol and following the same steps applied to the DNA fraction. The RNA pellet was resuspended in 12 μL of nuclease-free water. Each embryo was genotyped using its corresponding purified DNA fraction. The RNA samples were merged according to their genotype to generate from three to four biological replicates. Prior to library preparation, contaminating DNA remnants were degraded using the *TURBO DNA-free kit* (Ambion). Each RNA library was finally sequenced using an Illumina HiSeq 2500 system.

#### Downstream bioinformatic analysis

Reads were pre-processed trimming the Illumina universal adapters and the first 12 bases of each read to avoid k-mers using *Trimmomatic* (Bolger et al., 2014). Reads shorter than 50 bp and those with an average quality lower than 20 were filtered out (HEADCROP:12 MINLEN:50 AVGQUAL:20). Potential rRNA sequences were removed using *sortmerna* (Kopylova et al., 2012). Processed reads were then mapped against the last version of the medaka genome (ASM223467) using Hisat2 (Kim et al., 2019). Only reads with a high mapping quality (*samtools view -q 60*) were considered for further steps of the bioinformatics analysis. The software *htseq-count* was used to count the number of reads per gene (GTF from Ensembl version 99 was used as a reference). The subsequent analysis was performed using DEBrowser v1.14.2 (Kucukural et al., 2019). Genes with less than 10 reads on average were discarded (RowMeans < 10) and data were normalized following the relative log-expression (RLE) method. Potential batch effects were removed using ComBat. Differential gene expression analysis was carried out using the R package DESeq2 (padj < 0.05; -log FC = 1) https://www.r-project.org/. Raw and processed data were uploaded to the GEO database.

GO Terms were analyzed using GProfiler (Raudvere et al., 2019). The lists of *yap1* and *yap1b* DamID peaks obtained previously (Vazquez-Marin et al., 2019) were concatenated and, after assigning the closest gene using *Bedtools*, they were compared to the list of downregulated genes in our bulk RNA-seq to identify which genes are potential direct binding targets for YAP1 and YAP1B. As the Ensembl version used for the RNA-seq analysis is more recent that the one used for the DamID-seq analysis (Ensembl version 89), those genes which may have changed their identifier were not considered for this analysis. The statistical significance for the comparison between control and identified overlapping DamID peaks was calculated applying a two-proportion Z-test.

### Whole-mount in situ hybridization

cDNA from medaka embryos at stage 24 was used to amplify part of the coding sequence of medaka *no-tail* (ntl), *goosecoid* (gsc), *sox3, yap1* and *marcksl1b* genes. PCR products were cloned into pSC-A-amp/kan Strataclone plasmids (Agilent) to generate probes for wholemount in situ hybridization experiments (Table S1). Probes were synthesized using digoxigenin-11-UTP nucleotides (Roche) and the T3 or the T7 polymerase (Roche) depending on the insert orientation. In situ hybridization (ISH) was performed following a previous protocol (Thisse and Thisse, 2008). Medaka embryos at stage 15,16,17 and 18 were fixed in 4% PFA during 2 days, dehydrated in methanol, and stored at −20 °C.

Fluorescent *in situ* hybridization (FISH) were performed on medaka embryos at stage 16 using specific probes for *marcksl1b*. We followed the same protocol as for ISH, but with several differences from incubation with the anti-DIG antibody on: Incubation with blocking Buffer (2% Blocking Reagent from ROCHE in MABTween 1x) for 1 h, anti-digoxigenin-POD antibody (1:150 in Blocking Buffer) for at least 2 h at RT was used. The embryos were then washed six times with PBS 0.1% Tween at RT and then overnight at 4 °C. Later, the embryos were washed again with PBS-0.1% Tween and three times with Borate buffer (100mM Borate Buffer, 0.1%Tween), and stained with TSA amplification solution (50μg/mL TSA Fluorescein 5mg/mL in 100mM Borate Buffer, 0.1%Tw, 2% DS, 0.003% H2O2) for 1h at RT in the dark. Finally, embryos were incubated overnight at 4 °C with DAPI (Sigma) diluted 1:1000 in PBS-0.2% Tween. Stained embryos were mounted in FluoroDish plates as described previously. Confocal laser scanning microscopy was performed using an LSM 880 microscope (Zeiss). Only embryos that were mounted with their dorsal-anterior axis oriented in parallel to the cover glass bottom were used for the analysis. Images were processed using ImageJ (Schindelin et al., 2012).

**Table S1:**
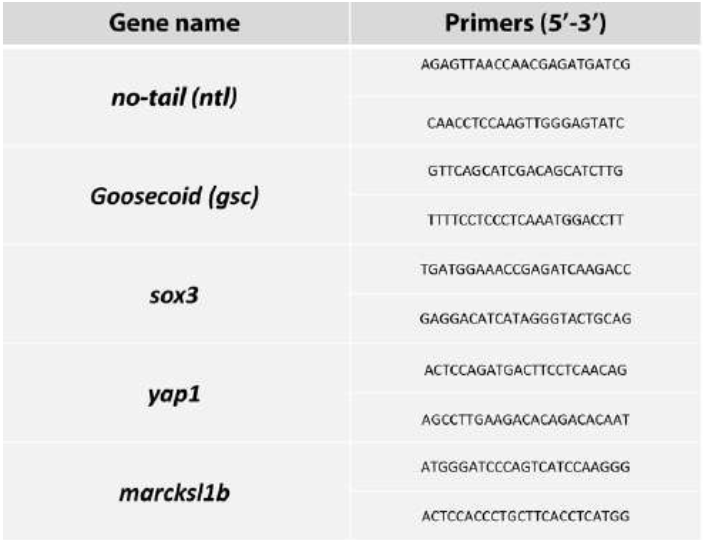
List of primers used to generate the RNA probes.

To establish a correlation between *marcksl1b* and cell density, we determined the centroid position of each nucleus, excluding the centroids that were closer than 8 pixels and those nuclei at the border of the image. Then, we calculated the mean of the distance between the five closest neighbors for each nucleus. Based on the *marcksl1b* expression pattern, we distinguished two different areas; an active area which shows a specific expression pattern for *marcksl1b* and an inactive area in which cells are not expressing *marcksl1b*. We represented the XY position of the nuclei’s centroids, with a circle or a triangle if they were localized in the active or the inactive area, respectively. The color gradient of each nucleus was dependent on the mean distance to its five closest neighbors. To analyze whether experimental groups were significantly different, two-sided Student’s t-tests were performed.

### mRNA Generation and injection

DNA plasmids containing, *yap1::mcherry, utrophin::GFP (given by Dr Norden), paxillin::mKate (Addgene 105974) and lynTdTomato* were linearized with NotI and then transcribed using the mMESSAGE mMACHINE SP6 Kit (Ambion) to synthesize capped mRNA. RNA was injected into one-cell-stage embryos (100-300 pg per embryo).

Embryos injected with *utrophin*::GFP and paxillin::*mKate* or *lynTdTomatoe* mRNA were fixed in PFA 4% at stage 16 for 2 days. Fixed embryos were washed with PBS-0.1% Tween and dechorionated with forceps. Finally, embryos were incubated overnight at 4 °C with DAPI (Sigma) diluted 1:1000 in PBS-0.1% Tween. Embryos were imaged with a Zeiss LSM 880 microscope (63x objective), taking images of a pixel size of 0.132 x 0.132 μm^2^ and a voxel depth 0.24 μm. Only embryos that were mounted with the dorsal-anterior axis oriented in parallel to the cover glass bottom were used for analysis. Imaged embryos were genotyped later to identify yap1/yap1b-related genotypes. Images were processed using ImageJ (Mahoney et al., 2005). For each image and channel, a maximum projection was generated. For XZ projections we used the Volume viewer plugin from ImageJ. To analyze the 3D morphology of the nuclei we applied fist a Gaussian Blur 3D filter (X, Y and Z sigma value was set in 2.0) to the blue channel. Then, we segmented and created a mask for these nuclei using the Watershed algorithm. To obtain the geometrical and morphological parameters we used the plugins 3D Geometrical measure and 3D shape measure from ImageJ. Downstream analyses were carried out using R. Nuclei smaller than 100 and bigger than 350 (volume unit) were excluded. To analyze whether experimental groups were significantly different, twosided Student’s t-tests were performed.

### Rockout treatment

Medaka embryos were dechorionated in vivo following a previous protocol (Porazinski et al., 2010) with minor modifications. First, embryos were rolled in sandpaper (2000 grit size, waterproof) to weaken their outer structure. Then, the embryos were incubated for 30 min at 28ºC in pronase at 20 mg/mL. Finally, after several washing steps, embryos were incubated for 60 min in hatching enzyme at 28ºC and transferred into a Petri dish with BSS 1x medium. Embryos at stage 15 were incubated with DMSO or Rho Kinase Inhibitor III (555553, Merck) at 250 μM dissolved in water for 2 h. Then embryos were fixed with PFA 4% for 2 days to perform the *in-situ* hybridization as explained previously.

## ACKNOWLEDGMENTS

We thank the CABD Proteomics, Aquatic Vertebrates and Functional Genomics facilities for their excellent technical assistance. This work was supported by grants awarded to JRMM from the Spanish Ministry of Science, Innovation and Universities: (References BFU2017-86339P,RED2018-102553-T, PID2020-112566GB-I00, and CEX2020-001088-M), and by the Marie Sklodowska-Curie H2020-MSCA-IF-2018-ST MechaPattern 834610 and La Caixa Junior Leader from Fundación Social La Caixa given to MAC.

---

**Supplementary Figure 1.**
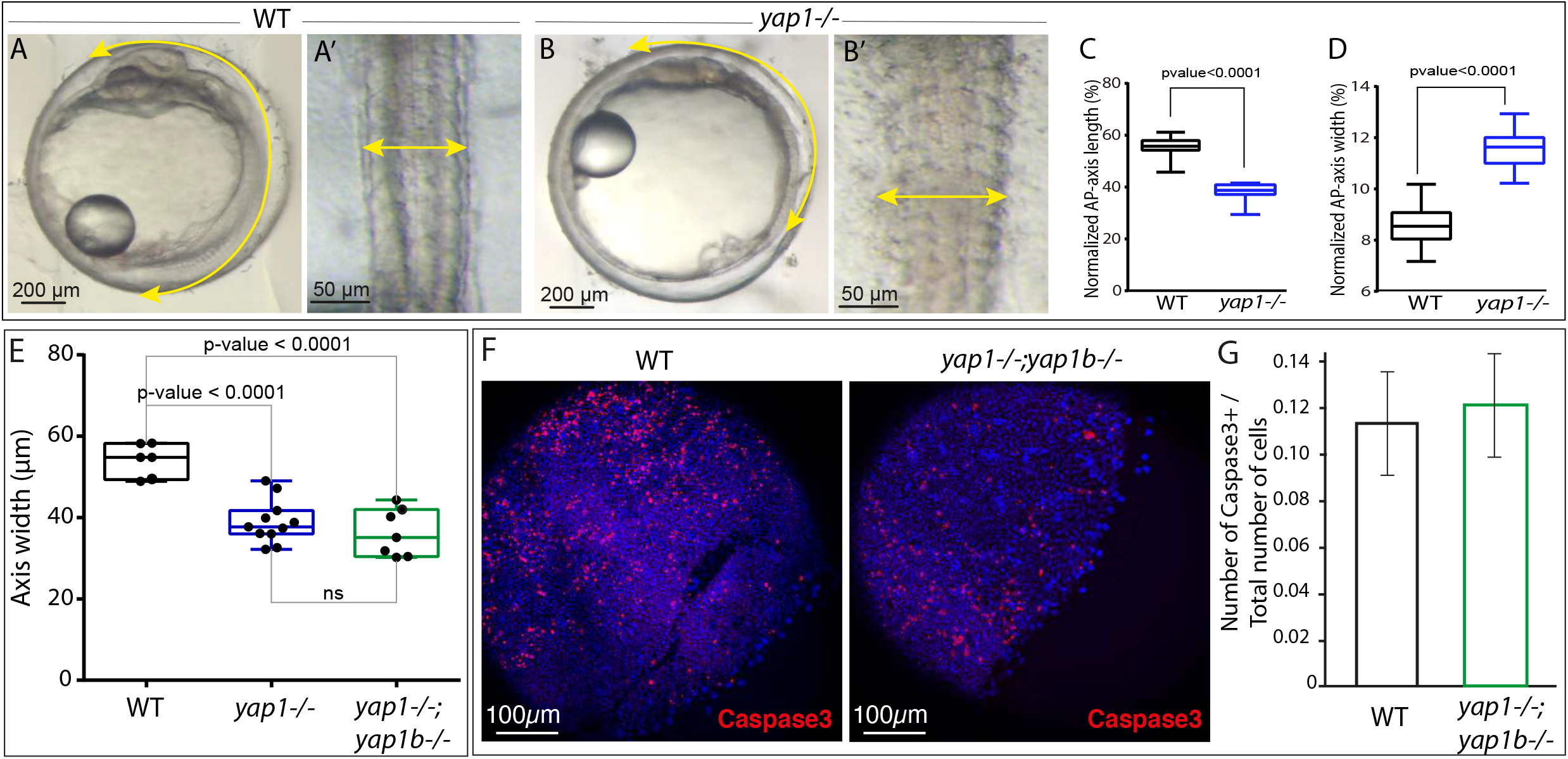
Analysis of *yap* mutants’ phenotype. **(A,B)** Brightfield images of WT and *yap1^-/-^* embryos at stage 24. Yellow double-headed arrows highlight the shortening and the widening of the A-P axis (A-B). A’, B’ correspond to magnifications of the axis. **(C,D)** Quantification of the normalized length (C) and width (D) of WT and yap1^-/-^embryo A-P axis. Boxes represent the quartiles; the whiskers indicate the maximum and minimum values. two-sided Student’s t-test was used to evaluate statistical significance. P-values < 0.0001. **(E)** Quantification of posterior axis width in WT, *yap1^-/-^* and *yap1^-/-^;yap1b^-/-^* embryos at stage 24. Boxes represent the quartiles; the whiskers indicate the maximum and minimum values. two-sided Student’s t-test was used to evaluate statistical significance. P-values < 0.0001. **(F)** Confocal images show Caspase 3+ cells in red and DAPI in blue in WT and *yap1^-/-^;yap1b^-/-^*. Scale bar 100 μm. **(G)** Quantification of caspase 3-positive cells per total number of cells in WT and *yap1^-/-^;yap1b^-/-^* embryos. The whiskers indicate the maximum and minimum values. Two-sided Student’s t-tests were performed to evaluate statistical significance. No significant differences were obtained. Scales bars 200 μm (A,B), 50 μm (A’-B’) and 100μm (F).

**Supplementary Figure 2.**
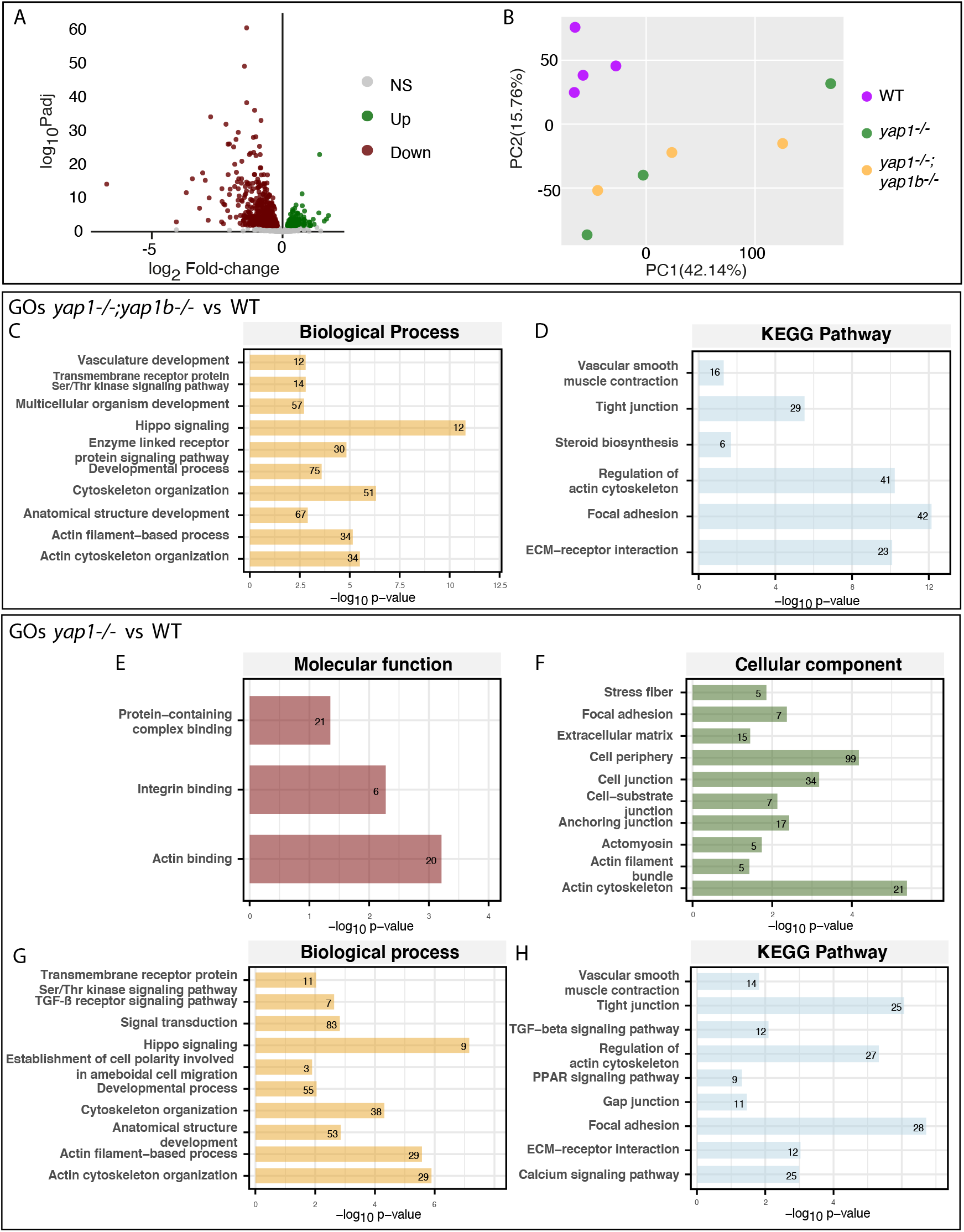
Yap-dependent transcriptional programs. Volcano plot graph showing the DEGs obtaining when comparing WT to *yap1^-/-^* embryos (A). Gray dots: no differentially expressed genes; Green dots: up-regulated genes in *yap1^-/-^* embryos compared with WT; Red dots: down-regulated genes in *yap1^-/-^* embryos compared with WT. Differential gene expression analysis was carried out using the R package DESeq2 (padj < 0.05; -log FC = 1). PCA graph showing the RNA-seq data variability between WT, *yap1^-/-^* and *yap1^-/-^;yap1b^-/-^* embryos (B). Gene Ontology (GO) enrichment of the DEGs in *yap1^-/-^;yap1b^-/-^* embryos compared with WT, classified in Biological processes (C) and KEGG Pathway (D). Gene Ontology (GO) enrichment of the DEGs in *yap1^-/-^* embryos compared with WT, classified in molecular function (E), cellular component (F), biological processes (G) and KEGG Pathway (H).

**Supplementary Figure 3.**
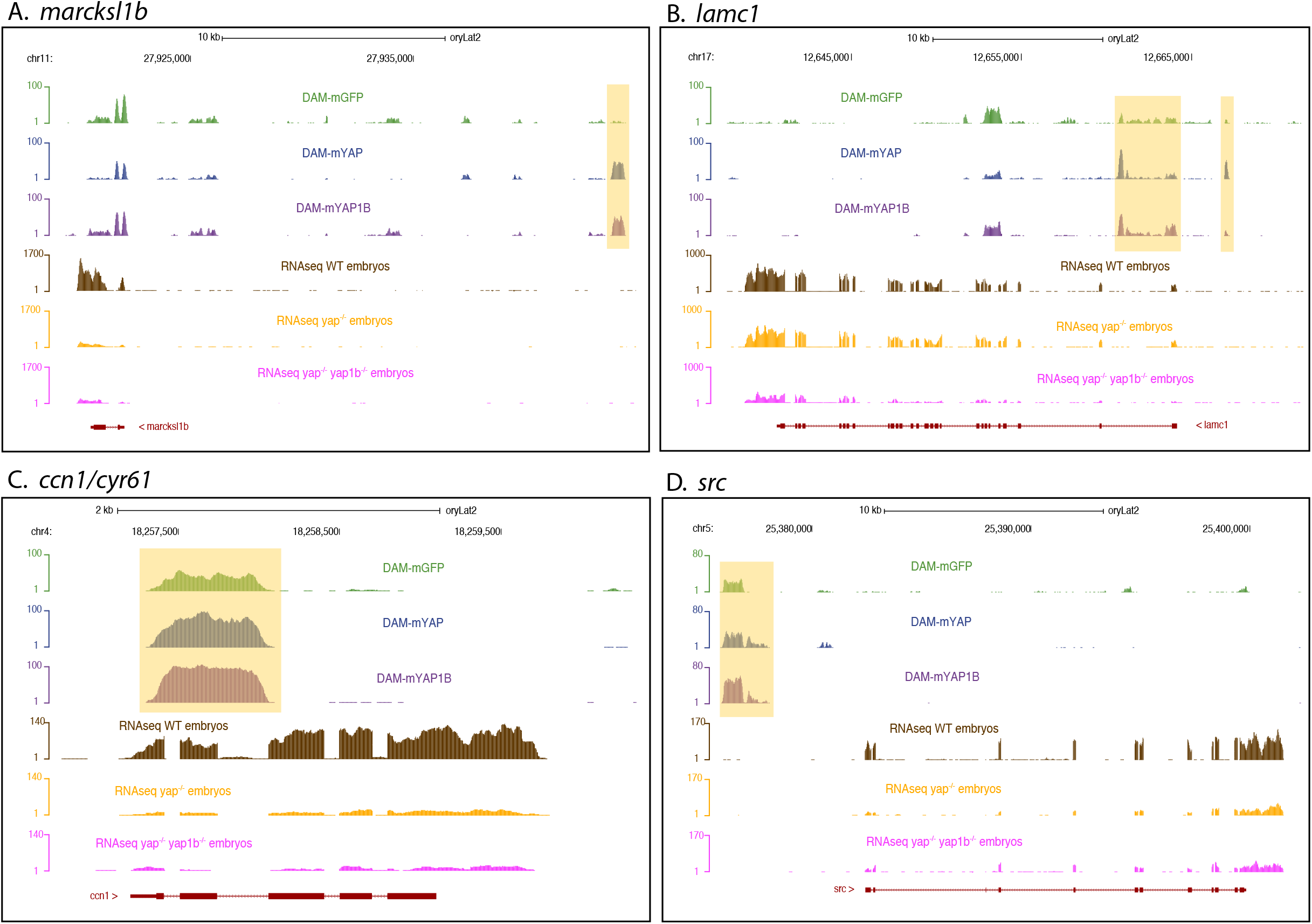
More significant DEGS’ DamID-seq and RNA-seq profiles. DamID-seq (Vázquez-Marín et al., 2019) and RNA-seq profiling of *Marcksl1b* (A), *lamc1* (B), ccn2/cyr61 (C) and src (D). The changes between the control, Yap1 and Yap1b in the DamID-seq peaks are highlighted in yellow.

**Supplementary Figure 4.**
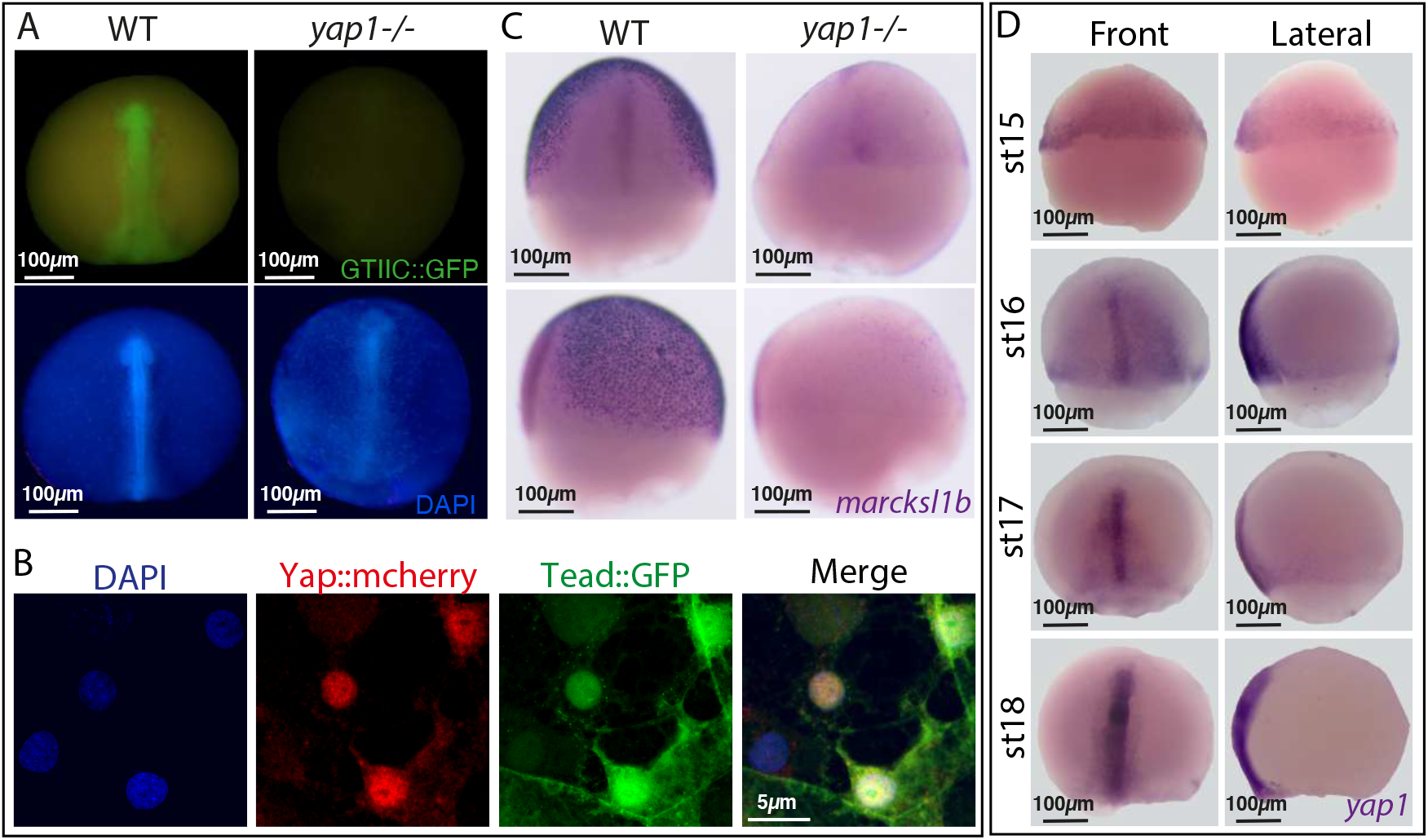
Yap activity dynamics during gastrulation. **(A)** GFP signal of WT and *yap1^-/-^ GTIIC::GFP* transgenic embryos at st 18. This later stage was used because it is when GFP signal become apparent under the fluorescent stereoscope. DAPI stainings of these embryos are also shown. Whole embryos are shown under the fluorescent stereo microscope. **(B)** DAPI counterstained confocal images of dorsal cells of *GTIIC::GFP* transgenic embryos injected with *Yap1*::mCherry. Blue channel: DAPI; Red channel: *Yap1*::mCherry signal; Green channel: GFP signal. **(C)** *In situ hybridization* analysis of the expression of *marcksl1b* in WT and *yap1^-/-^* embryos at st 16. Front and lateral images are shown. **(D)** *In situ hybridization* analysis of the expression of *yap1* in stages 15, 16, 17 and 18 embryos. Front and lateral images are shown. Scale bars 100 μm (A, C, D) and 5 μm (B).

Table 1: List of the most significant Differentially Expressed Genes (DEGs) of our RNAseq data (WT vs *yap1-/-;yap1b-/-*).

Movie 1: Tridimensional (3D) reconstruction of DAPI (blue) and Phalloidin (green) immunostained WT embryo posterior axis.

Movie 2: Tridimensional (3D) reconstruction of DAPI (blue) and Phalloidin (green) immunostained *yap1^-/-^* embryo posterior axis.

Movie 3: Tridimensional (3D) reconstruction of DAPI (blue) and Phalloidin (green) immunostained *yap1^-/-^;yap1b^-/-^* embryo posterior axis.

Movie 4: WT embryos injected with *Histone2b*::GFP for nuclei visualization were imaged in 4-min intervals over 8 hours. Manual cell tracking trajectories are represented with color lines.

Movie 5: *yap1^-/-^* embryos injected with *Histone2b*::GFP for nuclei visualization were imaged in 4-min intervals over 8 hours. Manual cell tracking trajectories are represented with color lines.

Movie 6: *yap1^-/-^;yap1b^-/-^* embryos injected with *Histone2b*::GFP for nuclei visualization were imaged in 4-min intervals over 8 hours. Manual cell tracking trajectories are represented with color lines.

Movie 7: Live confocal imaging of WT *GTIIC::GFP* transgenic embryos.

## Notes

### Competing Interest Statement

The authors have declared no competing interest.

